# DART: A GUI Pipeline for Aligning Histological Brain Sections to 3D Atlases and Automating Laser Microdissection

**DOI:** 10.1101/2025.09.03.673836

**Authors:** Rishi Koneru, Manjari Anant, Hyopil Kim, Julian Cheron, Justus M Kebschull

**Affiliations:** Department of Biomedical Engineering, Johns Hopkins University, Baltimore, MD 21205, USA; Department of Neuroscience, Johns Hopkins University, Baltimore, MD 21205, USA; Kavli Neuroscience Discovery Institute, Johns Hopkins University, Baltimore, MD 21205, USA; Center for Computational Biology, Johns Hopkins University, Baltimore, MD; Center for Functional Anatomy and Evolution, Johns Hopkins University, Baltimore, MD 21205, USA

## Abstract

The precise dissection of anatomically defined brain regions is the basis of many workflows in neurobiology. Traditionally, brain regions of interest are defined by visual inspection of tissue sections, followed by manual dissection. Recently, laser capture microscopes have been employed for more accurate dissection, but region identification is still challenging. This paper presents an open-source software pipeline DART (<u>D</u>issecting <u>A</u>tlas-<u>R</u>egistered <u>T</u>issue) that aligns histological brain sections to three-dimensional reference atlases and exports the resulting region-of-interest (ROI) contours for dissection by Leica Laser Microdissection (LMD) instruments. By integrating well-established image-processing libraries with a user-friendly graphical user interface, the software automates the traditionally time-consuming workflow of defining the boundaries of brain regions for dissection. With this pipeline, researchers can streamline tissue sampling for molecular analyses, while ensuring reproducibility and precision in ROI selection.

## Statement of Need

There is a growing demand for precise, automated methods to identify and dissect regions of interest (ROIs) in histological brain sections. The brain is a highly heterogeneous organ with distinct molecular and connectomic profiles across constituent regions, making accurate region dissection critical for meaningful biological interpretation. For example, proteomic analysis of brain regions and bulk RNA sequencing approaches rely on the dissection of histologically defined brain regions^1,2.^ Similarly, MAPseq and other barcoded connectomics techniques depend on the precise dissection of dozens to hundreds of brain regions to analyze how individual neurons project to these regions^3–6^. Laser microdissection (LMD) enables such precise dissection by cutting cell-stained tissue with an ultraviolet laser that can be controlled with high accuracy. However, current approaches using LMD generally rely on manual delineation of brain regions. This manual selection is error-prone and time consuming, especially when many samples need to be dissected. Moreover, the lack of a standardized region selection makes it difficult to ensure consistent, reproducible sampling across experiments and laboratories. There are many existing open-source tools that can align an image of a histological brain section to a reference brain atlas to identify and segment the brain regions (e.g. STalign, QuickNII, VisuAlign) ^7,8^. However, no tools exist that integrate atlas alignment of brain section images and brain region selection with Leica LMD software to enable the seamless and precise dissection of those regions.

Our open-source DART pipeline addresses this gap by providing:

1. Automated or semi-automated alignment of 2D brain sections to a 3D reference brain atlas (e.g., the Allen Brain Atlas CCFv3 2017^9^) and atlas-based segmentation.
2. Seamless integration with Leica LMD software for laser dissection.
3. Clear documentation and an accessible, modular codebase, facilitating further development and customization by the community.

Additionally, DART facilitates high-throughput processing of brain sections by minimizing user engagement time. The majority of the existing user engagement time is spent estimating and manually adjusting the atlas-to-brain section transform, which is facilitated by DART’s intuitive controls. The remaining steps require minimal input, allowing the user to attend to other tasks.

## Software Overview

### Key Features

1. **Brain Section Preprocessing**
  ∘ Automatic or user-assisted cropping and background intensity correction for raw histological images.
2. **Atlas Alignment**
  ∘ Combination of STalign^8^ for landmarked-based semi-automatic registration and Visualign^7^ for manual registration to a 3D reference brain atlas (e.g., Allen Mouse Brain Atlas^9^).
  ∘ Generation of affine and non-linear transformations to map 2D section coordinates onto known brain regions.
3. **ROI Definition and Export**
  ∘ User-guided selection of atlas-defined regions in the aligned sections.
  ∘ Conversion of these regions into outlines, polygons, or shape files suitable for use by a Leica LMD (xml files).
  ∘ Ability to store metadata (e.g., atlas region names) along with the exported ROI shapes.
4. **GUI Control**
  ∘ A GUI for real-time quality control and manual correction of section alignment.
5. **Modularity and Extensibility**
  ∘ Written primarily in Python, making use of libraries such as scikit-image, NumPy, and PyTorch for image processing and transformations.
  ∘ Open-source architecture allowing advanced users to plug in new registration algorithms, customize region-detection heuristics, or adapt to different species and atlases.
6. **High-throughput Multi-Sample Processing**
  ∘ Slide segmentation tools allow users to upload images of whole slides with multiple sections and export the ROI outlines.
7. **Integration with LMD**
  ∘ Simple shape import process using Leica’s.xml template.
  ∘ Flexibility in selecting the magnification at which laser dissection of ROIs is performed, with the ability to switch between magnifications as needed without requiring redrawing of shapes.

### Implementation and Architecture

1. **Programming Languages and Dependencies** The core modules are implemented in Python, leveraging libraries such as Tkinter, NumPy, Matplotlib, STalign, Shapely, NiBabel, and Pynrrd.
2. **Overall Workflow (Figure 1):**
  1. **Input:** The user provides high-resolution cell stained (e.g. DAPI or Nissl stained) images of the slide containing the brain section(s) (commonly in TIFF or PNG) and selects the corresponding section level in the 3D atlas.
  2. **Section Definition:** In the case that multiple sections are contained in a single image, the user delineates the bounding boxes for each section.
  3. **Run STalign:** A semi-automatic alignment between the atlas and the section(s) is performed using STalign.
  4. **VisuAlign Adjustment:** The user can visualize the atlas defined region boundaries on the cytoarchitecture to confirm alignment accuracy and, if necessary, manually adjust the alignment using VisuAlign.
  5. **ROI Selection:** The user selects ROIs (e.g., hippocampal subregions, cortical layers) to be exported. DART currently exploits the hierarchical region organization of the Allen Mouse Brain Reference Atlas to enable ROI definition of both coarse regions (e.g. primary motor area) and finer regions (e.g. layer 1 of primary motor cortex).
  6. **ROI Export:** These ROIs are then saved in an XML format that Leica LMD software can import, allowing precise laser dissections.
3. **Software Architecture:**
  ∘ DART follows an object-oriented framework (Figure 2)

**Figure 1.**
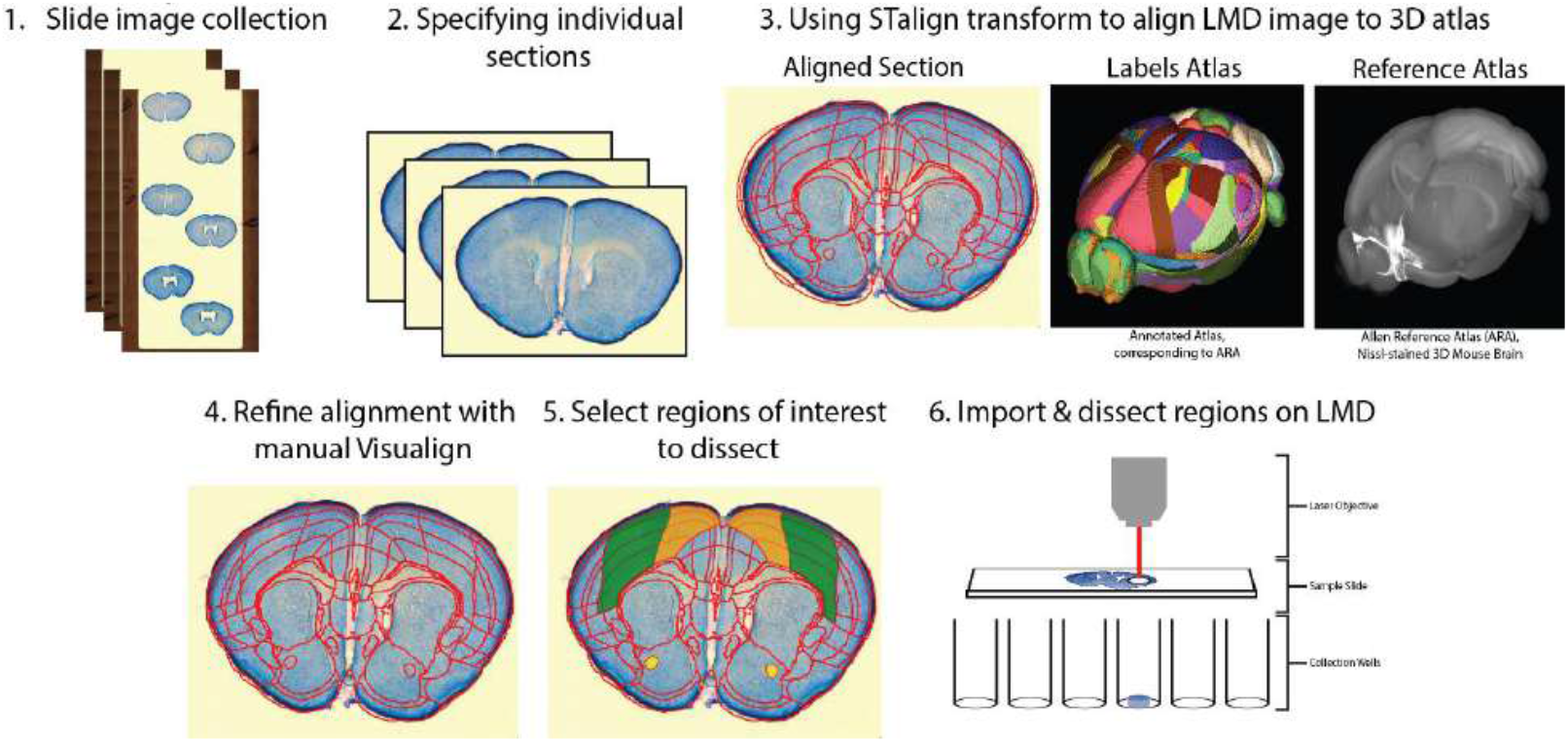
The DART workflow. The 3D renderings of the Allen atlas were produced using brainrender^9,10^. Cross Fiducial Example.

**Figure 2.**
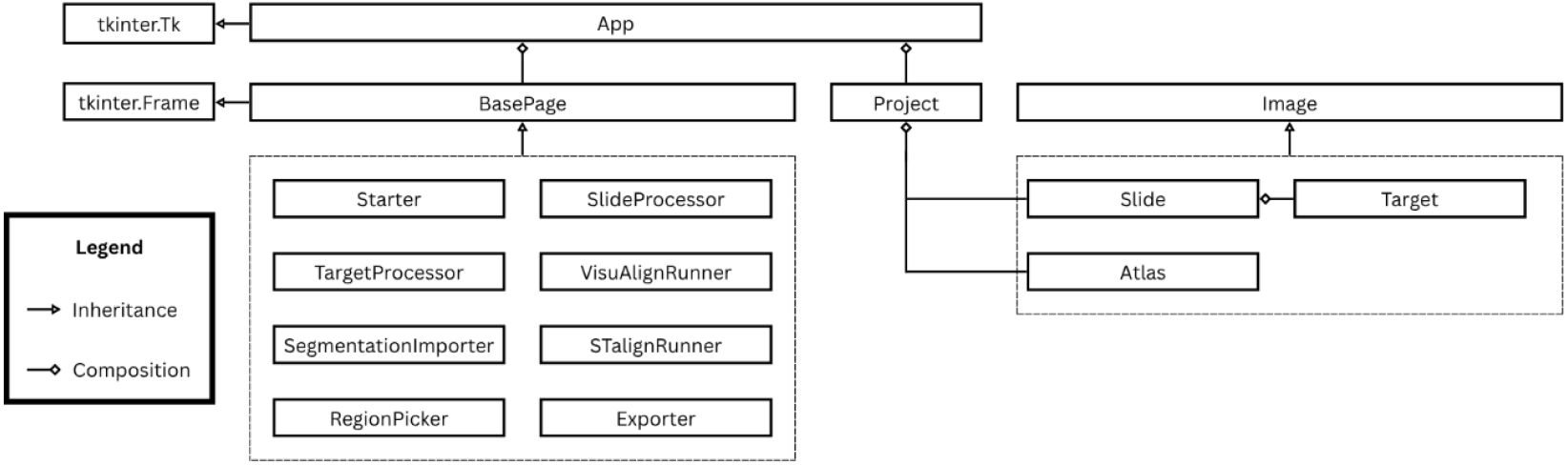
Class diagram of the object-oriented layout of DART

## Application

To demonstrate its utility, we applied DART to dissect the primary and secondary somatomotor areas and the anterior commissure from a coronal section of an adult mouse brain (Figure 3). Starting with a Nissl-stained section, we used DART to align the section to a DAPI-based atlas^11^ and generated outlines for the target regions. Importing these outlines into the Leica LMD system enabled precise laser cutting.

**Figure 3.**
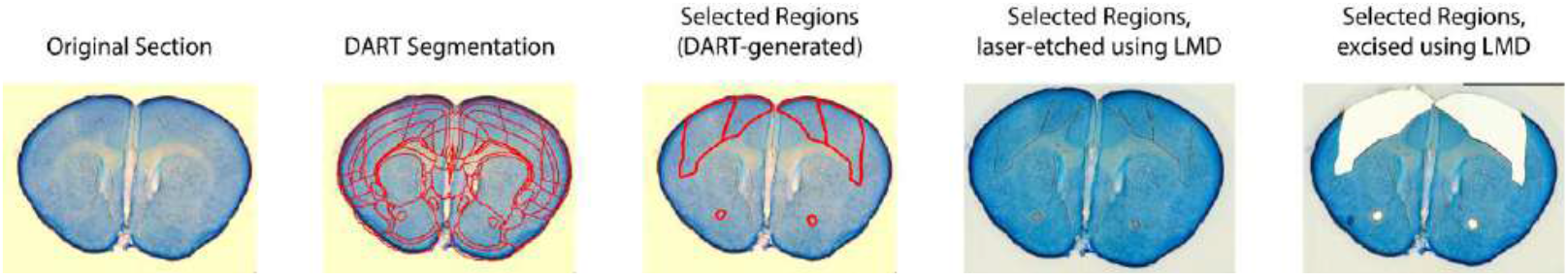
A DART use case. A DART application begins with an image of the original section. It then generates a segmentation, selects desired regions, and exports to the LMD for cutting or etching with the laser as desired. Note: Images were restitched using the Stitching plugin in ImageJ^12^.

## Discussion and Limitations

Our open-source workflow greatly simplifies consistent region identification and dissection across multiple brain sections. However, certain limitations remain:

- **Staining Variability**: Significant differences in staining intensity or section thickness can reduce alignment accuracy.
- **Atlas Mismatch**: Anatomical variability between individual subjects and the reference atlas may necessitate manual adjustment, especially in samples from disease models.
- **Tissue Damage**: Significant deformation to sections from holes or tears in the tissue can reduce alignment accuracy.
- **ROI Complexity**: Highly intricate or irregular regions might require manual editing of automatically generated ROIs.

Future improvements of DART may include support for alternative alignment algorithms, expanded atlas selection covering additional species and organ systems, and expanded compatibility with additional microscope systems beyond Leica LMD.

## Conclusions

Here we present an open-source pipeline that aligns histological brain sections to 3D reference atlases and seamlessly exports brain-region defined ROIs for Leica Laser Microdissection. By automating key steps in the tissue sampling process, researchers can reduce manual labor, improve reproducibility, and accelerate downstream molecular assays. The modular, documented codebase invites community contributions and adaptation to new species, atlases, and laboratory workflows.

## Code Availability

DART is hosted on Github: https://github.com/rk324/Dissecting-Atlas-Registered-Tissue

## Installation

DART is distributed as a pre-compiled Windows binary in a standalone folder that includes all necessary dependencies. To use the software, download and extract the entire folder from the [Google Drive link](https://drive.google.com/drive/folders/1OZ8UNjqNX_7eInwqjmnIVMsgPvRFYCfx?usp=drive_link), then run the main.exe file inside—no installation or separate Python environment is required. See the [tutorial](https://github.com/rk324/Dissecting-Atlas-Registered-Tissue/tree/main/tutorials) for instructions on usage.

## Data Collection

All mouse procedures were conducted in accordance with the Johns Hopkins University Animal Care and Use Committee (ACUC) protocols MO20M376 and MO23M346. Mice were maintained on a 12-hour light/dark cycle with ad libitum access to food and water. Brain sections for LMD were prepared as previously described^13^.

## Acknowledgments

We would like to thank the open-source community for providing foundational libraries such as VisuAlign, STalign, and Tkinter. This work was supported by SFARI grant 875575 and NIH grants DP1DA056668 and RF1AG078378 to JMK and a Kavli NDI Distinguished predoctoral fellowship to MA.

## User Guide

### Installation

`DART` is distributed as a pre-compiled Windows binary in a standalone folder that includes all necessary dependencies. To use the software, download and extract the entire folder from the Google Drive link, then run the `main.exe` file inside—no installation or separate Python environment is required.

### Getting Started

Prior to using `DART`, calibration points must be added to the slide. Although natural landmarks on the sample can be used, we recommend creating artificial calibration points that can be reliably located. To do this, use the laser to mark cross-shaped fiducials in the slide membrane at the top left, top right, and bottom right corners of the slide (**Figure 1**). Then, take an overview image of the whole slide, including fiducial crosses, to use in `DART`.

**Figure 1.**
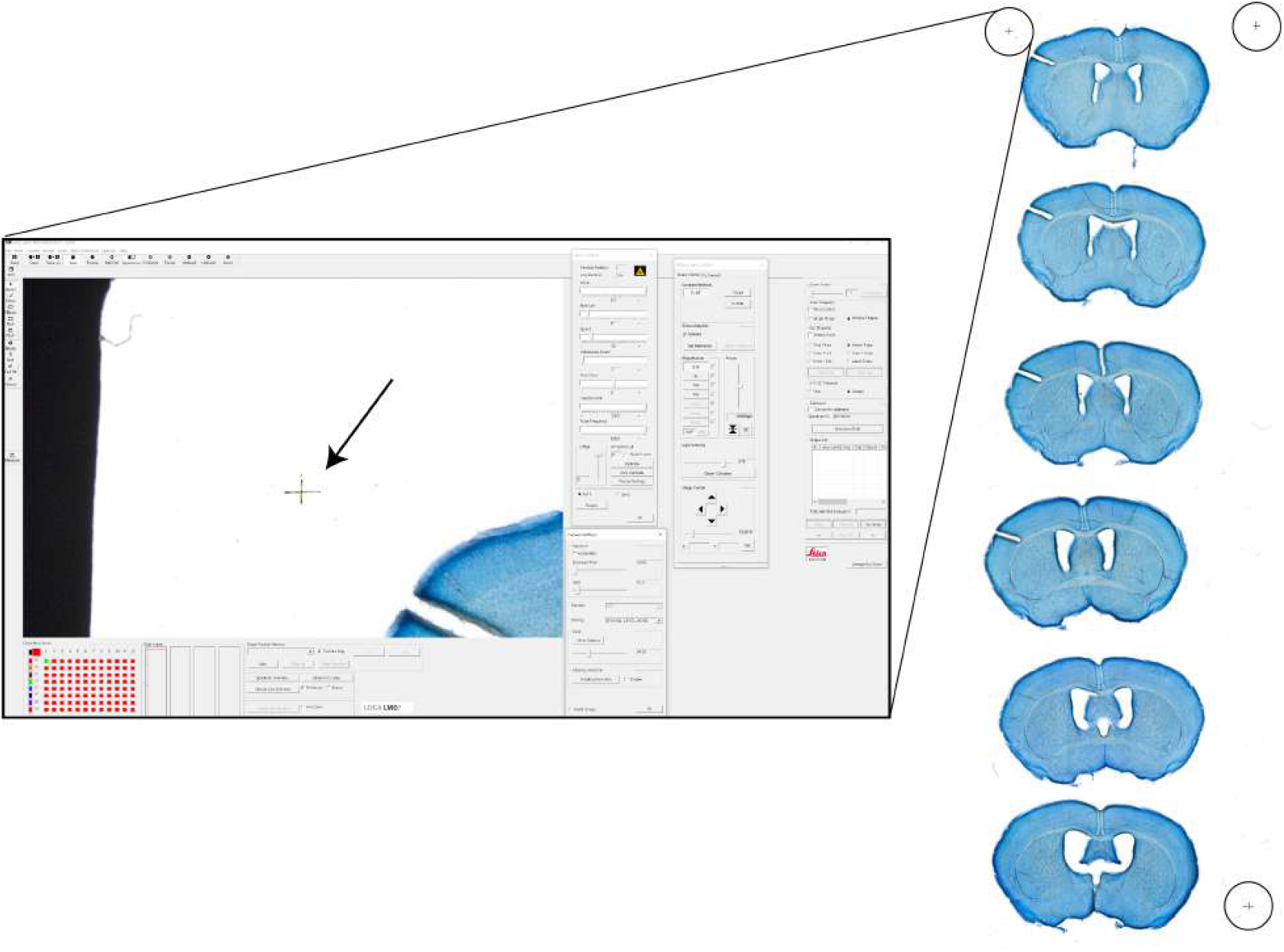
Cross Fiducial Example

### Load the Data

On the `DART` starter page (**Figure 2**), select an atlas from several options with varying resolutions and imaging workups. Atlases comprise three components: a reference atlas that contains the spatial cell density of the organ of interest, a labels atlas that maps each voxel from the reference atlas to an ID corresponding to a specific region in the tissue, and a table containing information about the regions and their hierarchical structure. In addition to selecting an atlas, select the sample images by clicking **Browse** and selecting the folder containing the images. `DART` will load each image in the folder as a separate slide and create a subfolder to store results and intermediate files. The name of the subfolder will follow the format `DART-[datetime]`.

**Figure 2.**
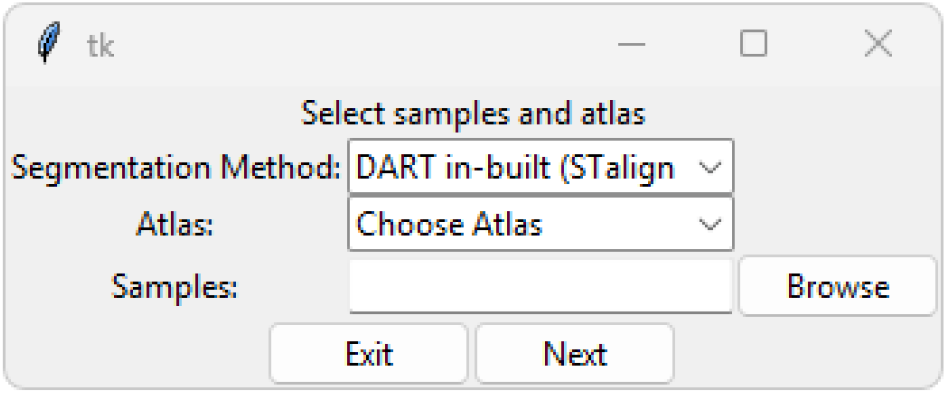
Starter Page

### Mark Calibration Points and Select Sections

On the slide processing page (**Figure 3**), the **Add Calibration Points** tab is selected by default. Annotate the calibration points (marked earlier with the LMD) by clicking on the calbration point and then selecting **Add Point** on the top panel. Next, switch to the **Select Slices** tab on the top panel and drag a box around each section to delineate the individual sections. Since multiple sections can be mounted on a single slide, this allows for bulk processing of several sections on each slide. When annotating images in this software, a common color scheme is used. Red annotations have not been saved, green annotations have been saved, and orange annotations have been recently saved and can be removed with a corresponding button at the top of the page. This color scheme is also applied when annotating calibration points, and it continues throughout the software.

**Figure 3.**
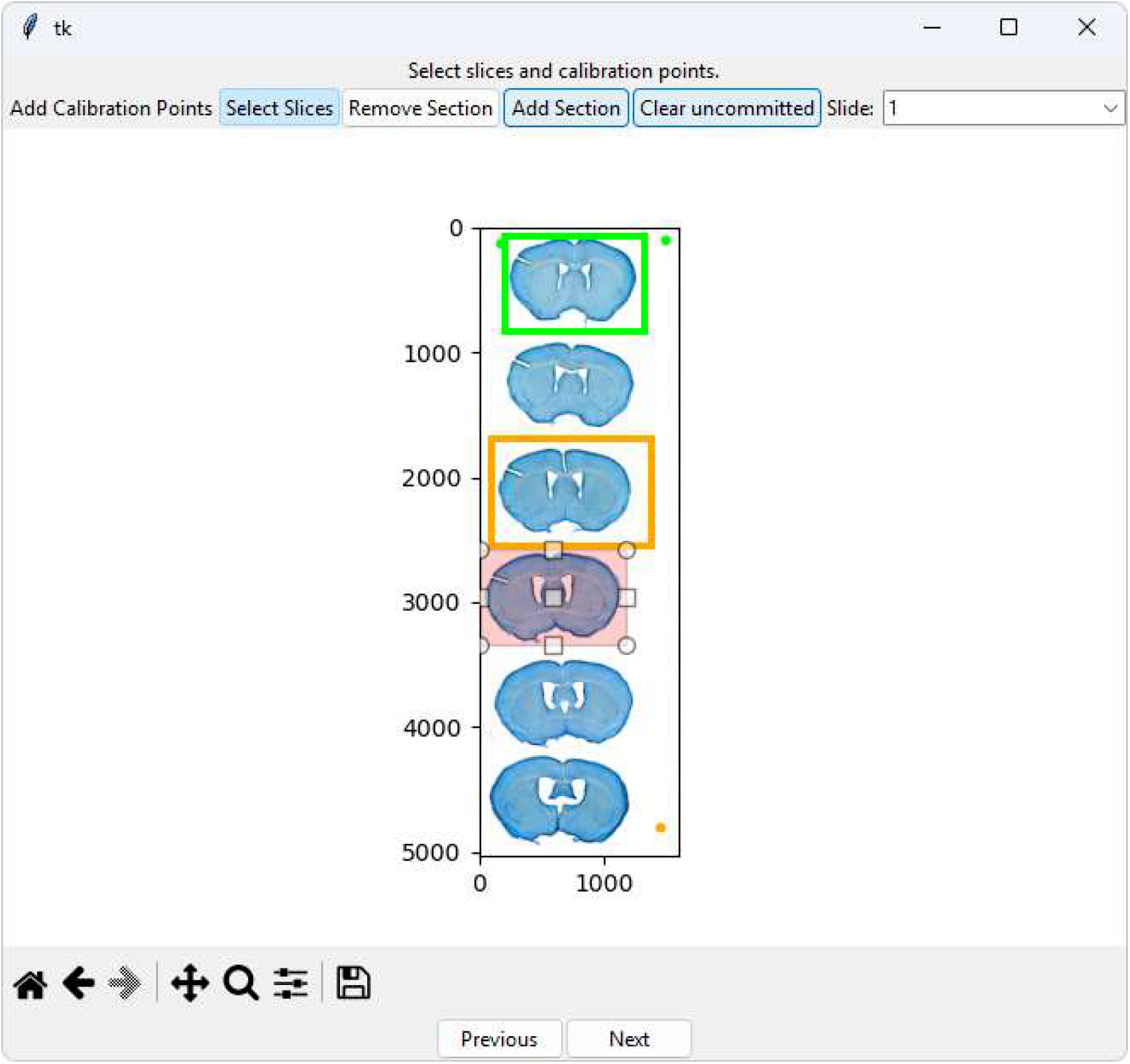
Slide Processing Page

### Prepare for STalign

The Section Processor Page (**Figure 4**) serves three functions: estimation of an affine transformation to map the atlas to the target section, annotation of landmark points, and adjustment of STalign parameters. Use the bottom slider to approximate the section in your 3D atlas that best matches the section. Use the three sliders on the right to adjust the rotation angles of the atlas to more closely match the slice. This is your initial “guess” for the alignment, and even if it’s not perfect, it will greatly speed up on converging on a proper alignment.

**Figure 4.**
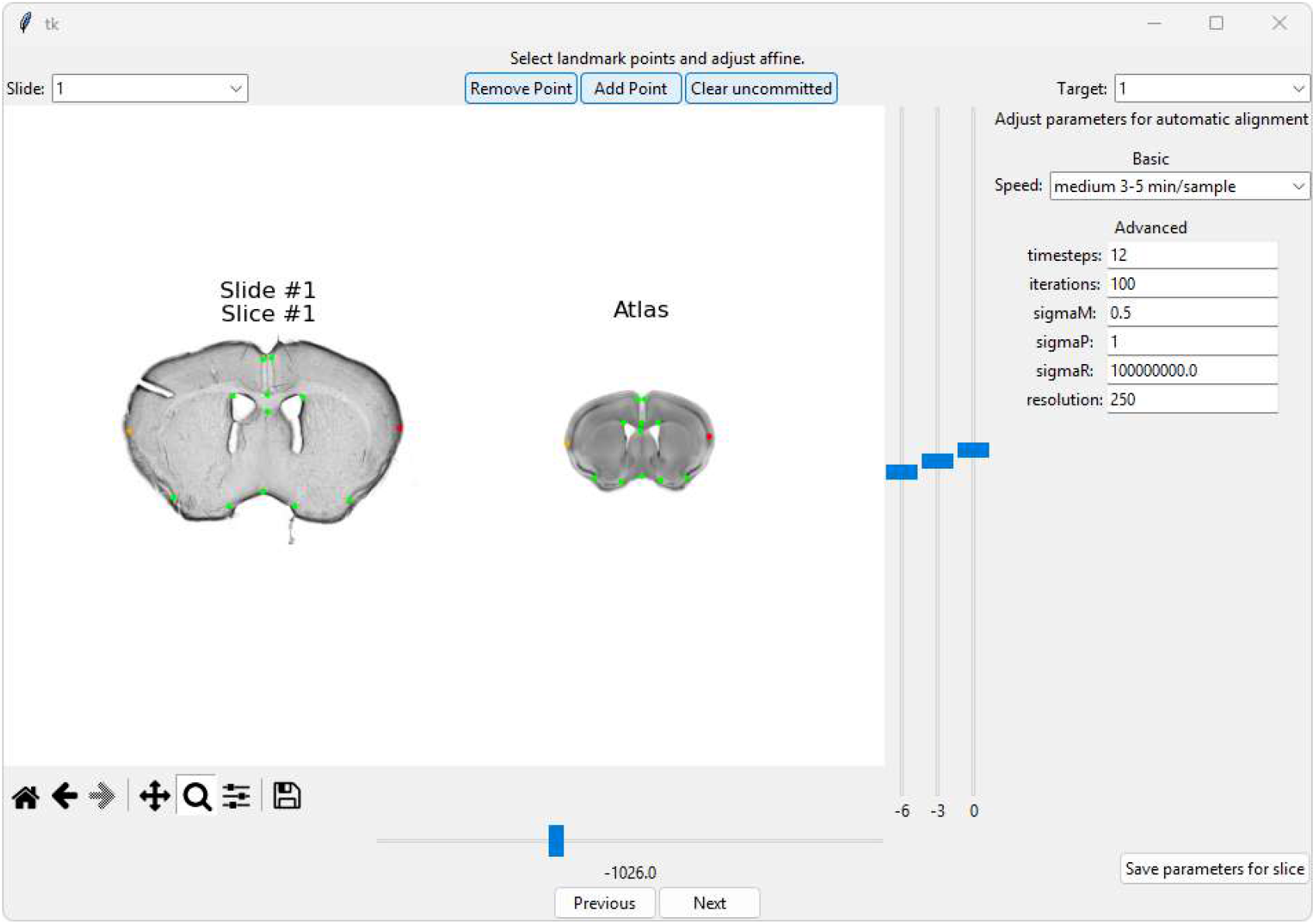
Section Processor Page

Next, add landmark points by clicking corresponding landmarks on the target section image and the atlas image, then clicking “Add Point” in the top panel.

Finally, you can tune the parameters of STalign either at a high-level through the dropdown menu or in detail through the text entries. The slower transforms are generally more accurate. You can save these parameters by clicking **Save parameters for slice** in the bottom right.

If you have multiple slices, you can navigate to the different slices using the **Target** panel on the top right, and initialize an alignment for each one.

### Run STalign and View Results

In the STalign Runner Page (**Figures 5, 6, 7**), STalign is run on the section images when the “Run” button is clicked. Upon completion of STalign, the results are displayed by overlaying the calculated region boundaries over the section image (**Figure 7**).

**Figure 5.**
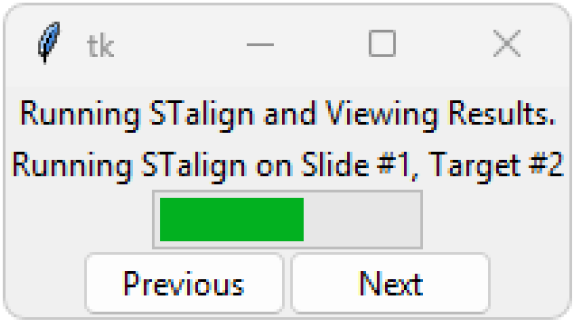
STalign Runner Page

**Figure 6.**
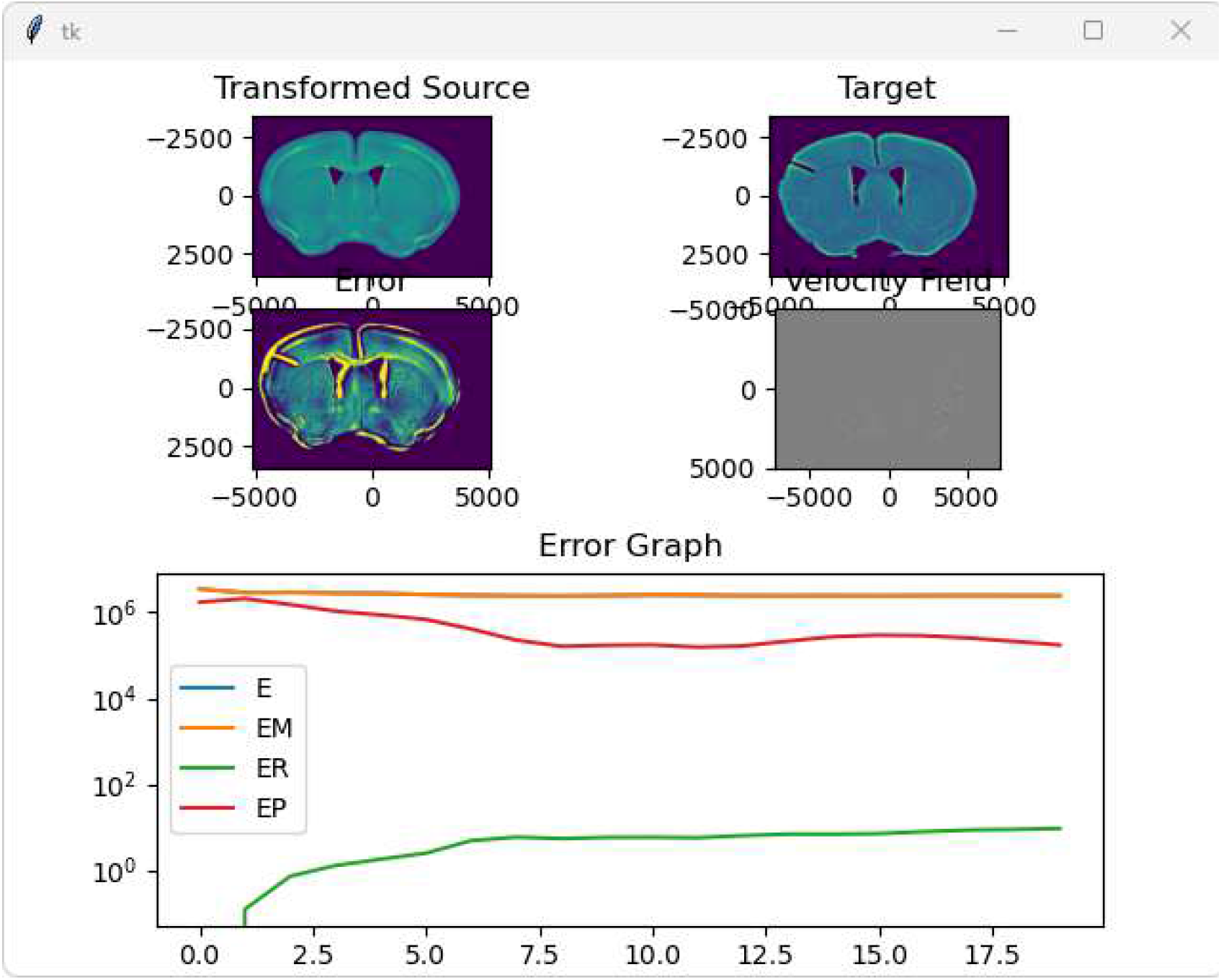
STalign Progress Monitor

**Figure 7.**
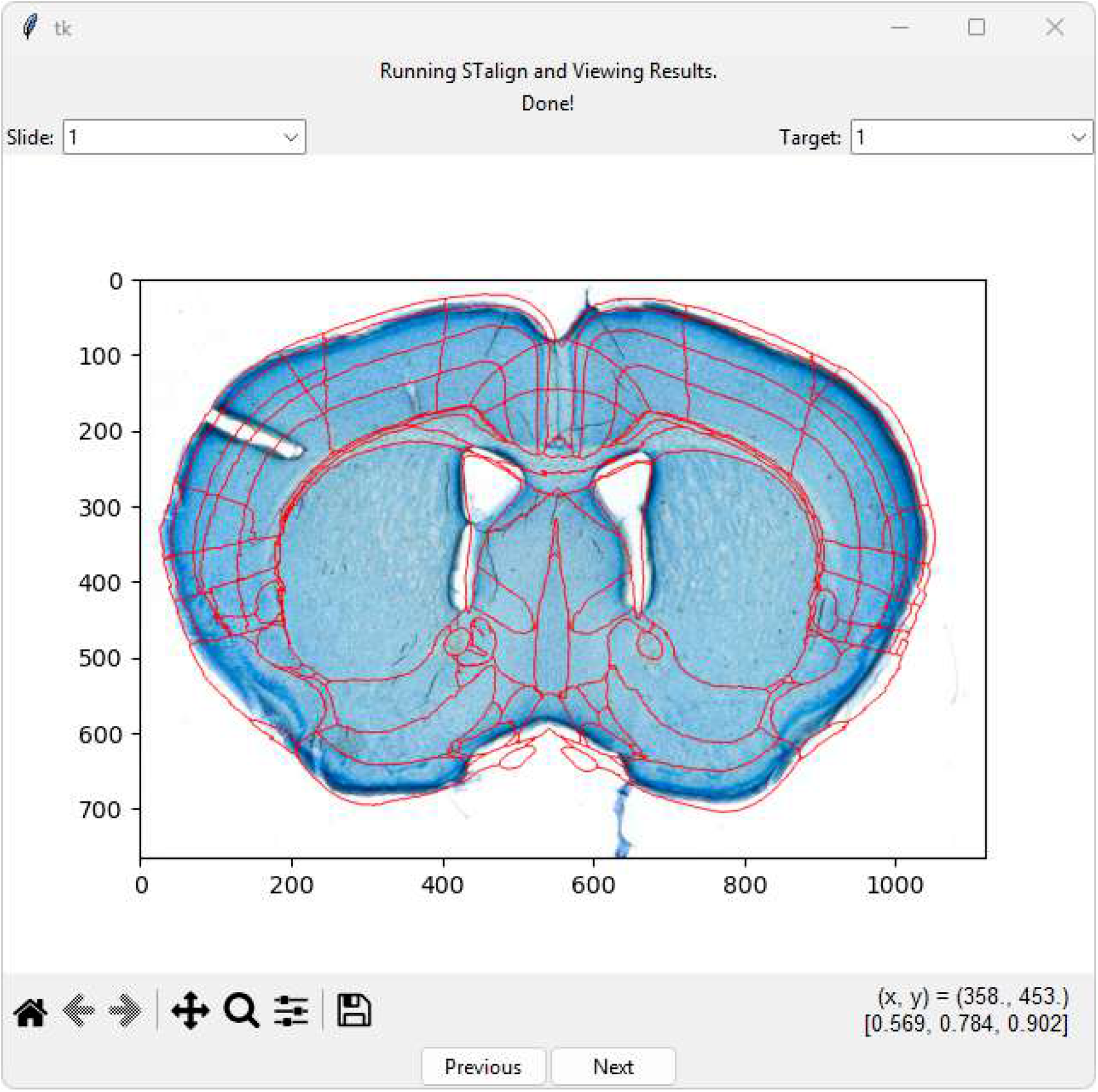
STalign Results Display

### Adjust Alignment

In the VisuAlign Runner page (**Figure 8**), make manual adjustments to the alignment using VisuAlign (**Figure 9**). This enables greater control over the alignment. Since VisuAlign is a separate software, `DART` opens it through the command terminal, when you click **Open VisuAlign**.

**Figure 8.**
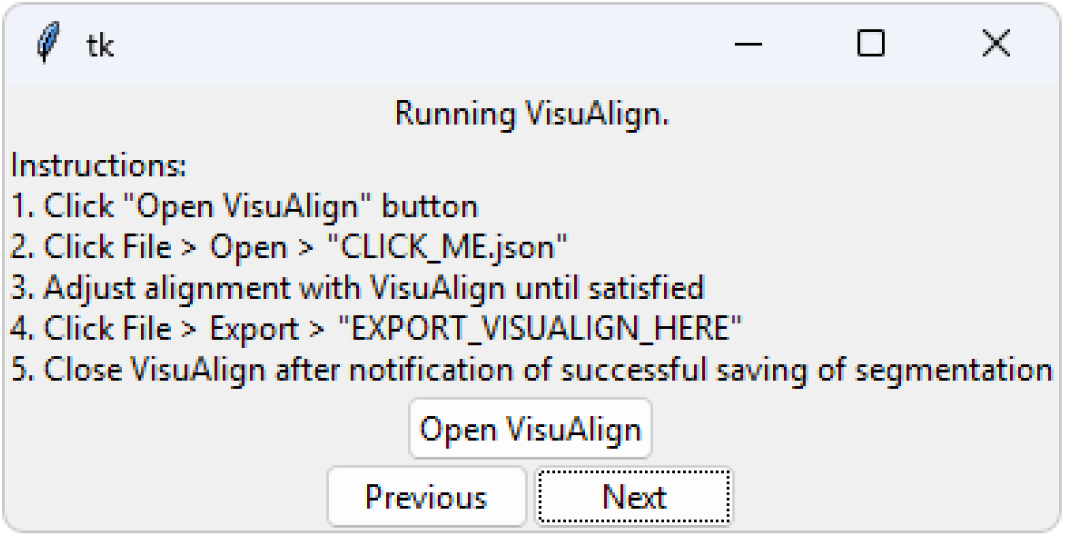
VisuAlign Runner Page

**Figure 9.**
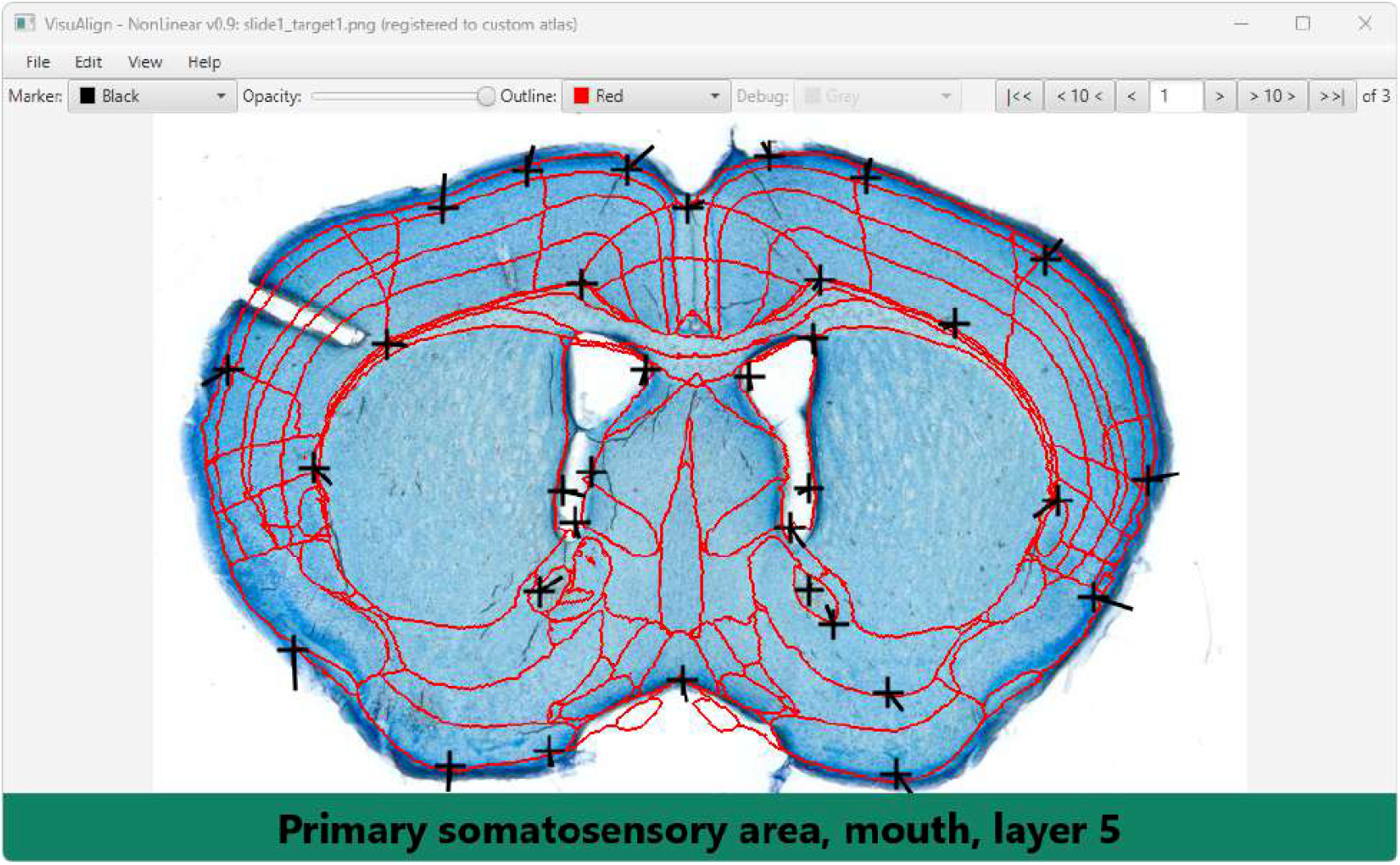
VisuAlign

You will find the `CLICK ME` file in the `DART-[datetime]` subfolder with your section images. You can do this by hitting the space bar on a region you wish to move, which should make a cross appear. You can then move the cross around to adjust the local alignment. If you have multiple slices, you can navigate slices with the arrows in the top right corner. After adjusting the alignment, export the alignment to the EXPORT_VISUALIGN_HERE folder, and close VisuAlign.

### Select ROIs

In the region selection page (**Figure 10**), select the regions of interest (ROIs). This can be done by either clicking on the ROIs in the image or by navigating to and toggling the checkbox of the region in the tree view. Each has three possible states: unchecked, checked (marked by a check mark), and tristate (marked by a filled box). Unchecked regions will not be exported for dissection. Checked regions and all their child regions are stitched together for combined dissection. Tristate regions are the ancestors of checked regions that are exported and dissected separately. This distinction between checked and tristate regions allows dissection of fine and broad groups of regions. For example you could define and dissect the entire cerebral cortex by using a checked box for parent **Cerebral Cortex** and all child regions. Or, you could use a filled box (tristate) for parent **Cerebral cortex**, and all immediate child regions (i.e. Visual Cortex, Somatosensory cortex, etc) that are checked are exported and dissected separately.

**Figure 10.**
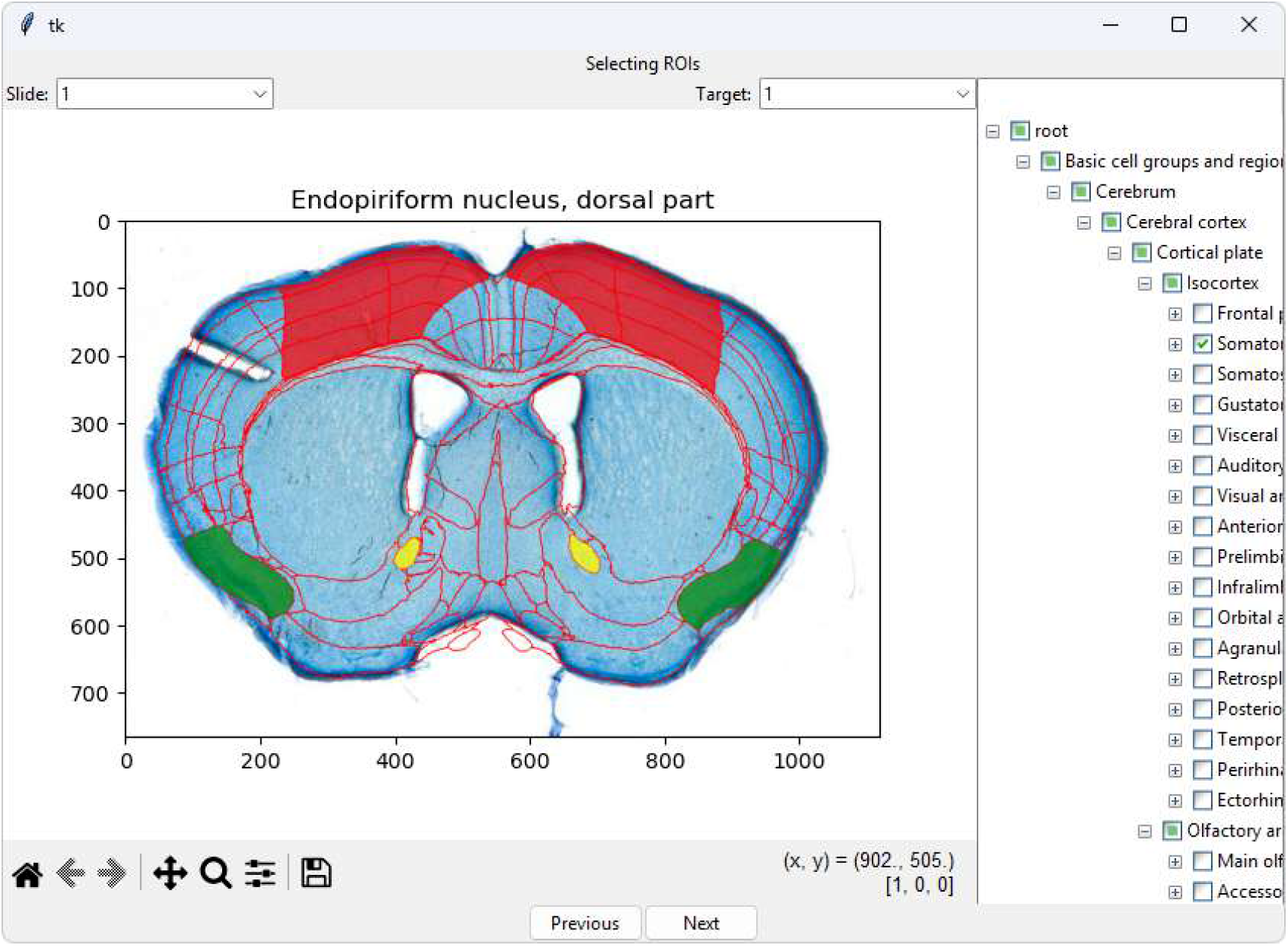
Region Selection Page

### Export ROI Boundaries

The export page (**Figure 11**) allows the user to select sections for export. All the sections of a slide can be exported together in one batch file for the slide, or individual sections can be exported. This allows the user to group their LMD cutting jobs as desired. The `.xml` files necessary for LMD dissection are in the `DART-[datetime]` subfolder, under the subfolder `output`.

**Figure 11.**
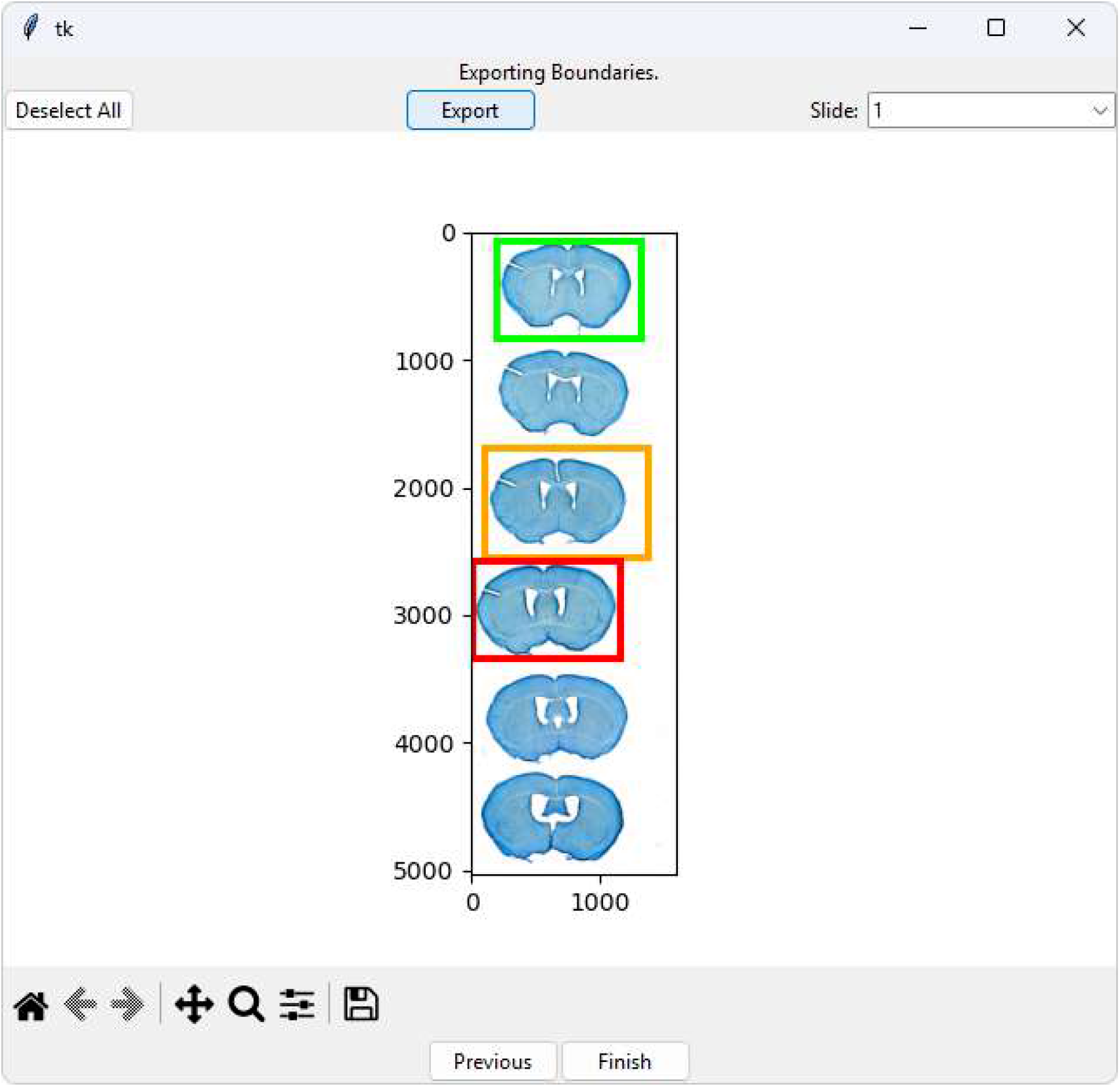
Export Page

### Import to LMD

Before importing shapes onto the LMD, switch the LMD to the desired objective for cutting. For example, if the user wants to cut their shapes using the 10x objective, then they must first switch to that objective. Then, click File > Import Shapes and select the `.xml` file containing the shape(s) to be imported. This will trigger a series of prompts to select shapes and calibrate the LMD. Navigate through these prompts until the final prompt, “Use the actual magnification for all imported shapes?”, is reached. Click “Yes”.

The shapes list should populate with the imported shapes. Select a shape to view it overlaid on the section, and click “Start cut” to initiate the laser dissection process for this shape. Alternatively, all the shapes may be selected and dissected. Note that `DART` automatically assigns wells to each shape, spaced out with one well in between. These well assignments may be adjusted in the Leica LMD software.

